# A Practical and Cost-Effective Approach to Long-Fragment eDNA Sequencing for High-Resolution Genetic Diversity Assessment

**DOI:** 10.1101/2025.10.11.681776

**Authors:** Satsuki Tsuji, Naoki Shibata, Tetsu Yatsuyanagi, Yusuke Fuke

## Abstract

Environmental DNA (eDNA) analysis is increasingly recognised as a valuable method for assessing genetic diversity. However, its resolution and applicability are limited by the short length of sequences that can be analysed (typically < 400 bp) and high analytical costs. This study developed a practical, low-cost long-fragment eDNA analysis method using commercial full-length plasmid sequencing via a nanopore platform and evaluated its effectiveness in assessing population genetic structure. 1 L of surface water was collected from 52 sites across Hokkaido, Japan, targeting *Barbatula oreas*. Two mitochondrial regions (ND5 and cyt *b*; approximately 1,000 bp each) were species-specifically amplified, circularised, and sequenced. Library preparation took 2.5 hours, with a total cost per sample of 4,390 JPY (≈25.55 EUR, ≈29.87 USD). High-quality reads were obtained from 34 samples, allowing for the reconstruction of multiple haplotypes per region through haplotype phasing. The eDNA concentration required to achieve a 50% sequencing success was within a range easily attainable for common species.

Phylogenetic analysis using 62 concatenated haplotypes (1,968 bp) obtained from each sample identified two clades and multiple regional subgroups, providing higher-resolution phylogeographic information than the previous study. Furthermore, the differentiation of each clade and group was suggested to reflect geological and climatic events. These results demonstrate the feasibility and utility of long-fragment eDNA analysis for evaluating genetic diversity, and its broad application is anticipated in ecological research, conservation management, and environmental policy formulation.

## 1 Introduction

Monitoring and conservation of biodiversity are critical global priorities, and environmental DNA (eDNA) analysis has been increasingly recognised as an innovative approach to complement or replace conventional field sampling methods (Liu et al., 2025; Rodríguez-Ezpeleta et al., 2021; Rourke et al., 2022). Since the first report of eDNA detection in macroorganisms in 2008 (Ficetola et al., 2008), technological advancements have progressed rapidly, driven in part by the widespread availability of high-throughput sequencing technologies (Tsuji et al., 2019; Valentini et al., 2016; Yao et al., 2022). Among various eDNA approaches, eDNA metabarcoding, which amplifies DNA fragments from multiple taxa using universal primers and assigns obtained sequences to species, has established itself as a method for efficiently assessing species biodiversity (Miya et al., 2020; M. Zhang et al., 2023a). This technique is well-suited for both short- and long-term monitoring and large-scale biodiversity surveys (Daraghmeh et al., 2025).

More recently, not only species-level detection but also eDNA-based inference of intraspecific genetic variation has emerged as a promising approach in fields such as phylogeography, conservation genetics, population genetics, and landscape genetics (Andres et al., 2023; Couton et al., 2023; Sigsgaard et al., 2020).

In the evaluation of intraspecific genetic variation based on eDNA analysis, methods such as amplicon sequencing of short DNA fragments, target/non-target shotgun sequencing, target capture-based methods have been used (Andres et al., 2021; De Barba et al., 2024; Jensen et al., 2021; Manel et al., 2025; Sigsgaard et al., 2016; Tsuji et al., 2020a). These methods have successfully detected the genetic diversity of the target species, but each has several limitations.

Amplicon sequencing of short DNA fragments (typically < 400 bp) often has low sequence variants, limiting the detection and estimation accuracy of genetic differences within and between populations, as well as the application of traditional population genetics tools/indicators (Nakajima and Tsuri, 2024; Tsuji et al., 2023b). While other methods could obtain more comprehensive genetic data, they require samples with relatively high eDNA concentrations of the target species or low contamination from non-target species (ex., samples obtained from footprints, tank water or high-density habitats) (Andres et al., 2023; De Barba et al., 2024; Manel et al., 2025; Pinfield et al., 2019). Additionally, both the required volume of eDNA sample and the per-sample analysis cost are several to several tens of times higher compared to the amplicon sequencing method (Jensen et al., 2021). Furthermore, target capture-based methods are still in the early stages of application, and several challenges remain for their practical implementation, including technical complexity in both molecular and bioinformatic workflows, low and biased sequencing coverage, and difficulties in ensuring reproducibility (Hintikka et al., 2022; Jensen et al., 2021; Suchan et al., 2016). Therefore, the development of eDNA-based methods that allow simpler and more cost-effective assessments of genetic diversity from environmental samples at a highly practical level is expected to contribute broadly to ecological research, conservation management, and environmental policy formulation.

Long-read amplicon sequencing analysis, which has undergone rapid technological advancement and increasing societal adoption in recent years, has the potential to enable the simple and cost-effective analysis of variations in longer eDNA fragments amplified by PCR (Doorenspleet et al., 2025; Maggini et al., 2024a). Although long-read amplicon sequencing was initially limited by relatively high read error rates (5–10%), recent improvements in flow cells, duplex sequence technique and bioinformatics tools have substantially improved its accuracy and practicality (Jain et al., 2015; Ni et al., 2023; T. Zhang et al., 2023b). Furthermore, commercial long-read amplicon sequencing services are now available from many companies worldwide (Nakagawa et al., 2025). These services allow long-read sequencing to be conducted with reduced initial cost and setup effort. Furthermore, Nakagawa et al. (2025) proposed a cost-effective and streamlined method in which long PCR amplicons are circularised using T4 ligase and sequenced using a full-length plasmid sequencing service via the nanopore platform. Full-length plasmid sequencing service using Nanopore technology is now offered by several internationally established companies [e.g., Eurofins Genomics (US), AZENTA Life Sciences (US), Plasmidsaurus (US)], typically available at less than one-tenth the cost of standard commercial long-read amplicon sequencing services. Therefore, by employing a full-length plasmid sequencing service, it is expected that the approximately 1000-bp eDNA fragments typically targeted in tissue-based surveys can be detected from environmental samples in a simple and low-cost manner.

To verify the utility of a full-length plasmid sequencing service for eDNA-based intraspecific genetic analysis, it is necessary to empirically evaluate whether it can detect known genetic variation and improve the resolution of population genetic inference compared to conventional methods. In this study, the Siberian stone loach *Barbatula oreas* (or *B. toni* as a synonym; Fig.1A; Dyldin et al., 2023; Kottelat, 2012), widely distributed in Hokkaido, the northernmost part of Japan, was selected as the model species. This species completes its life cycle in freshwater (Hosoya, 2019) and is suggested to have expanded its range under the influence of geological and climatic events such as land-bridge formations, orogenic activity, and glacial-interglacial cycles (Ooyagi et al., 2018; Yatsuyanagi et al., 2024). These processes should have shaped complex patterns of genetic structure both within and among their regional populations. Yatsuyanagi et al. (2024), using short-read eDNA metabarcoding, have revealed two major genetically distinct clades in Hokkaido, each comprising three subpopulations. Given these characteristics and background, *B. oreas* provides an ideal system for examining the extent to which genetic differences can be detected with high accuracy using eDNA-based population genetic analysis employing full-length plasmid sequencing. Furthermore, clarifying the genetic structure of this species at high resolution presents a valuable opportunity to understand how biodiversity in Hokkaido has been shaped by geohistorical and climatic changes.

**Figure 1.**
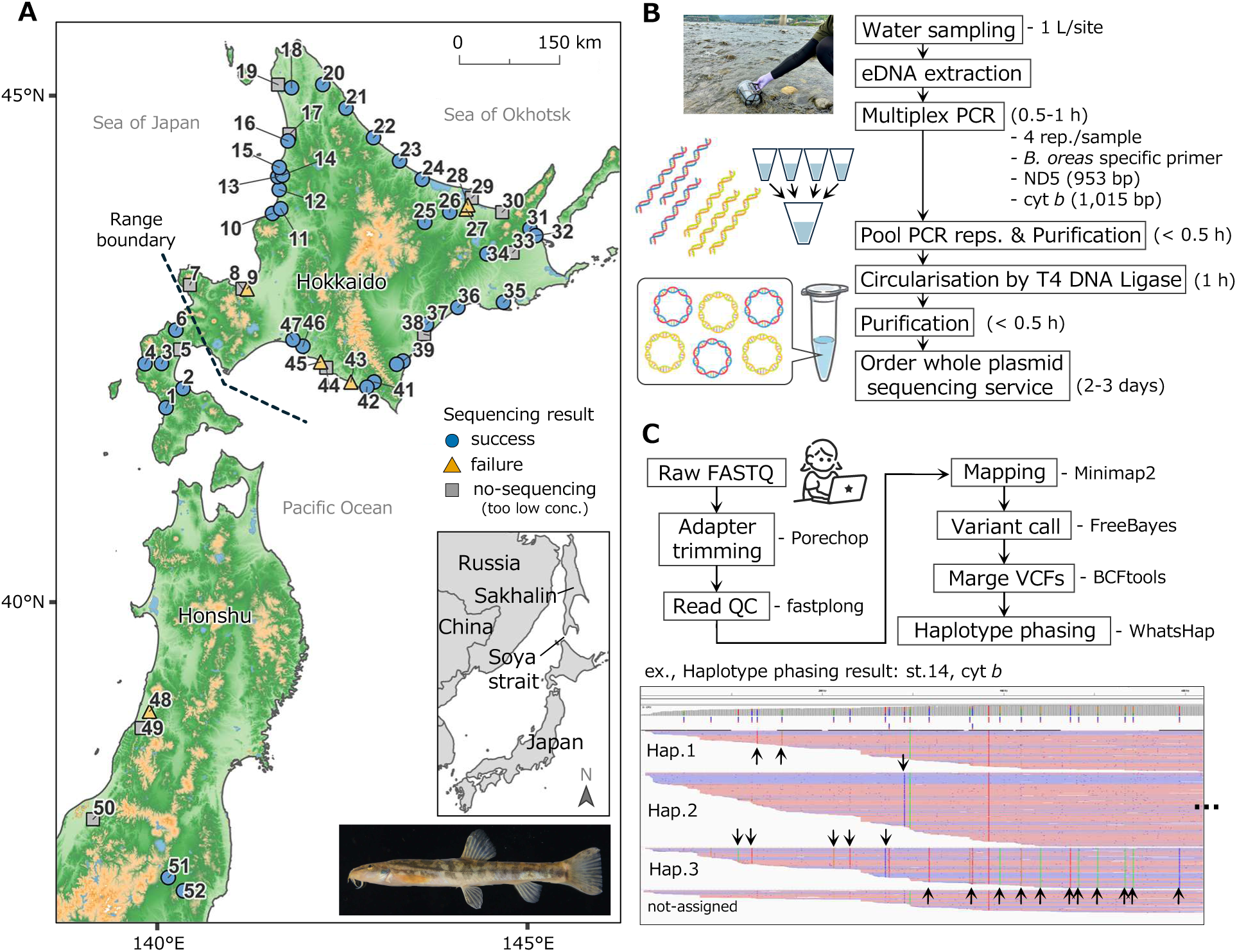
(A) *Barbatura oreas* and the overview of sampling sites. The black dotted line indicates the distribution boundary of *B. oreas*. Point shapes and colours indicate the result of long-fragment eDNA sequencing. (B) Workflow for the library preparation and estimated hands-on time. (C) Outline of bioinformatics analysis workflow. The lower panel shows an example of a screen when the haplotagged BAM file is visualised using IGV (Integrative Genomics Viewer) (ex. st.14, cyt *b*). The variant sites for each haplotype are indicated by arrows

The purpose of this study is to develop a simple and cost-effective long-fragment eDNA analysis method and to demonstrate its effectiveness as a method for evaluating genetic diversity. To achieve this, 1 L of surface water was collected from 52 sites across Hokkaido, Japan, targeting *B. oreas*. Two mitochondrial regions (approximately 1,000 bp each) were species-specifically amplified. For sequencing, a simple and low-cost protocol was established that combines the circularisation of long eDNA fragments and a commercial full-length plasmid sequencing service. Based on the co-occurrence patterns of SNPs in each long-read, haplotypes present in each sample were reconstructed. Phylogeographic and population genetic analyses were performed to investigate the population genetic structure of *B. oreas* in Hokkaido.

Furthermore, the concentrations of target eDNA in each sample and of the prepared libraries, both of which were required for successful sequencing, were examined. Accordingly, we discuss the usefulness, potential and future challenges of our proposed long eDNA analysis method for evaluating genetic diversity.

## 2 Materials and methods

### 2.1 Study sites and eDNA collection

Water sampling for eDNA collection was conducted on a total of 52 sites (Fig. 1). Most of the samples were collected in September 2024, and some were collected in August 2024 or August 2023 (Table S1). At each sampling site, 1 L of surface water was collected using a disposable plastic cup, and benzalkonium chloride (0.5 mL, 10% w/v; Fujifilm Wako Pure Chemical Corporation, Osaka, Japan) was added to preserve eDNA (Yamanaka et al., 2017). The water sample was vacuum-filtered on-site using a GF/F glass fibre filter (diameter: 47 mm, mesh size: 0.7 μm; GE Healthcare Japan, Tokyo, Japan) and filter holder (PP-47; ADVANTEC). After filtration, filter samples were folded in half, wrapped in aluminium foil, and immediately frozen in dry ice.

DNA extraction from filter samples was performed according to Tsuji et al. (2025b). Briefly, each filter sample was placed in the upper part of the empty spin column (EconoSpin, EP-31201; GeneDesign, Inc., Japan) and pre-centrifuged at 5,000 g for 1min to minimise the inhibition by remaining water, including benzalkonium chloride (Tsuji et al., 2022). The excess water that fell into the 2 mL tube was discarded. The filter was incubated with a mixture consisting of 200 μL ultrapure water, 250 μL Buffer AL, and 25 μL proteinase K for 40 min at 56 °C. Following the incubation, the spin columns were centrifuged at 6000 g for 1 min. Then, 500 μL of 99.9% ethanol was added to the collected liquid and mixed well by pipetting. The DNA solution was purified using a DNeasy mini spin column (Qiagen, Germany) according to the manufacturer’s protocol. Finally, the DNA was eluted in 100 μL of Buffer AE. The Buffer AL, proteinase K and Buffer AE were used as supplied with the DNeasy Blood and Tissue kit. In all molecular experiments, filter pipette tips were used to avoid contamination. The eDNA samples were stored at − 25 °C until DNA library preparation.

### 2.2 Species-specific primer set development for Nanopore sequencing

The mitochondrial NADH dehydrogenase subunit 5 (ND5) and cytochrome *b* (cyt *b*) sequences of *Barbatula oreas* and its closely related species inhabiting Hokkaido Island and Tohoku district (*Lefua nikkonis*, *L. echigonia*, *Cobitis* sp. BIWAE type C, *Misgurnus chipisaniensis*, *M. anguillicaudatus, Niwaella delicata*) were downloaded from the National Centre for Biotechnology Information database (NCBI; https://www.ncbi.nlm.nih.gov/) (Table S2). However, quantitative fish eDNA metabarcoding suggests that these closely related species are rarely found at the survey sites in this study (also see Results, Table S13). The obtained sequences were aligned using MAFFT (https://mafft.cbrc.jp/alignment/server/), and primers were manually designed at positions containing 1 to 3 species-specific nucleotide variations for *B. oreas* within the five bases from the 3′ end: Bto_Cytb-F (5′-[PHO]-TGA AAC TTC GGA TCA CTC CTG G-3′), Bto_Cytb-R (5′-[PHO]-TAC TAG GGC AAG CTC ATT CTA GC-3′); Bto_ND5-F (5′-[PHO]-GTC ACC AAC TGG CAC TGG A-3′), Bto_ND5-R (5′-[PHO]-CAT CTT TTG AAA AGA AGC CTG CAA G-3′). The primer was synthesised with a 5′-phosphate modification (5′-phosphorylated). An in silico test was conducted using Primer-BLAST (https://www.ncbi.nlm.nih.gov/tools/primer-blast/) with default settings to evaluate the species specificity of the designed primer set.

### 2.3 Library preparation for long-read amplicon sequencing

An overview of the library preparation workflow is presented in Fig. 1B. To avoid contamination, the laboratory was completely separated before and after the PCR and the experimenter was not allowed to return to the pre-PCR room on the same day. The ND5 (953 bp) and cyt *b* (1,015 bp) of *B. oreas* were amplified by multiplex PCR (four technical replicates per sample). The multiplex PCR was performed in a 20-µL total volume of reaction mixture containing 10.0 µL of 2 × KOD One (TOYOBO, Osaka, Japan), 0.6 µL of forward and reverse primers (10 µM; Bto_Cytb-F/R, Bto_ND5-F/R), 5.6 µL of sterilised distilled H2O, and 2.0 µL eDNA template.

KOD One is known for its high accuracy (approximately 80 times that of Taq DNA polymerase), amplification efficiency (1–10 kb: 5 sec./kb), and resistance to crude samples (https://x.gd/TRldt2). To avoid PCR inhibition, 12 µL of each eDNA sample was purified in advance with ×1.2 Sera-Mag SpeedBeads Carboxylate-Modified Magnetic Particles (Hydrophobic) (Cytiva, MA, USA), eluted in 12 µL of TE buffer (pH 8.0), and used for PCR. The thermal conditions were 10 s at 98°C, 5 s at 55°C, 10 s at 68°C, repeated 43 cycles.

Pooled PCR replicates for each sample were purified with × 0.7 Sera-Mag SpeedBeads and finally eluted in 12 µL of low TE buffer. Subsequently, 1.2 µL of T4 DNA ligase buffer (Nippon Gene, Tokyo, Japan) and 0.5 µL of T4 DNA ligase were added and incubated for 60 min at 16°C to circularise target fragments. After incubation, the circularised DNA was purified with ×1 Sera-Mag SpeedBeads and eluted in 13 µL of low TE buffer. Following instructions from the analysis company, the DNA library obtained at high concentrations was diluted to 50 ng/µL using low TE buffer. Since DNA concentrations that are too low may prevent sequencing, samples were selected for analysis using a threshold of approximately 25–30 ng/µL concentration (Table S1).

Full-length plasmid sequence analysis using the nanopore platform was performed by the Plasmid-EZ service (GENEWIZ, from AZENTA Life Science, Tokyo, Japan).

### 2.4 Bioinformatics analysis

An overview of the bioinformatics analysis workflow is presented in Fig. 1C. Raw Nanopore reads were first trimmed to remove adapter sequences using Porechop (v. 0.2.4), with default settings. Quality filtering of reads was subsequently performed using fastplong (v. 0.3.0), with a minimum phred quality score threshold of 30. Only high-quality reads were retained for downstream analysis. Filtered reads were mapped separately to reference sequences for the ND5 (LC894542) and cyt *b* (AB100917) using Minimap2 (v. 2.30) with the map-ont preset. Read group information was added during mapping, and sambamba (v. 1.0.1) was used to convert, sort, and mark duplicates in the resulting BAM files. Variant calling was conducted using FreeBayes (v. 1.3.10) across multiple assumed ploidy levels (two–six), with the following options: --pooled-continuous, --haplotype length 500, --min-coverage 5 and --min-alternate-fraction of 0.20.

Variant files were merged across samples for each gene and ploidy level using BCFtools merge command (v. 1.22). Phasing of variants was performed using WhatsHap (v.2.8), assuming several ploidy levels (two to six), to generate phased VCFs. Consensus haplotype sequences were generated using BCFtools consensus command, specifying individual samples, ploidy level, haplotype number, and the phased VCFs. This step produced haplotype-resolved consensus FASTA sequences for each sample and gene region. In parallel, WhatsHap haplotag command was used to assign haplotype tags (HP:i:N) to individual reads within each BAM file based on the phased VCFs. Following this, WhatsHap haplotagphase command was applied to refine haplotype phasing using both the phased VCF and haplotagged BAM files, resulting in sample-specific rephased VCFs. Finally, haplotype frequency estimation was performed using SAMtools view command. For each haplotagged BAM file, the total number of reads and the number of reads assigned to each haplotype tag were counted. These counts were compiled into haplotype frequency tables for each sample, gene region, and ploidy level.

### 2.5 Haplotype identification and sequence pair concatenation

To identify the haplotype for each target region in each sample, a threshold of 20% for variant allele frequency (VAF) was set, and the number of variant sites per genetic region was determined (Table S1). Note that VAF refers to the frequency of each base at the corresponding position in the mapped read, not mismatches with the reference sequence (ex., Table S3). Based on the number of variant sites, the number of haplotypes and their sequences were determined (Table S1): (1) No variant sites: 1 haplotype (the consensus sequence obtained from the BAM file), (2) One variant site: two haplotypes (obtained from the consensus sequence, each differing only at the single variant site), (3) two or more variant sites: based on the haplotype frequencies estimated by WhatsHap at each ploidy level, the most frequent ploidy and its estimated haplotype sequence were selected. However, if the frequencies were similar between ploidies, the haplotagged BAM file was visually confirmed using IGV (Integrative Genomics Viewer) to determine the ploidy to use. To use longer sequences for analysis, sequence pairs between two target regions in each sample were concatenated. The sequence pairs were selected based on the detection frequency of each haplotype and its regional subgroup, prioritising combinations that were as consistent as possible (Table S4, S5, Fig S1).

### 2.6 Statistical analyses

All statistical analyses and graphic illustrations were carried out using R software v. 4.3.2 (R Core Team, 2023), and statistical values were evaluated at a significance level of α = 0.05. To identify the concentration conditions of eDNA samples required for applying our proposed sequencing method, the relationship between the number of *B. oreas* eDNA copies in the samples before PCR (12S rRNA, ca. 170 bp; quantified by the quantitative fish eDNA metabarcoding described below) and the concentration of the final sequencing library (including circularised 953 bp of ND5 and 1,015 bp of cyt *b*) was examined using Spearman’s rank correlation test. The effect of eDNA concentration and library concentration on the probability of long-read amplicon sequencing success was examined using a generalised linear model (GLM) with a logit link function. We defined “sequencing success” as cases where valid data were obtained for both ND5 and cyt *b*. Using the obtained model, the eDNA concentration and library concentration required for 50% and 100% probability of sequencing success were calculated, respectively.

For each target region, to test for significant genetic divergence among clades and subgroups, a hierarchical analysis of molecular variance (AMOVA) was performed using the amova() function of the ‘pegas’ package (Paradis, 2010). Pairwise genetic distances among haplotypes were calculated using the Kimura 2-parameter (K80) model (Kimura, 1980). Significance was assessed after 1,000 permutations.

### 2.7 Phylogenetic analyses

For each target region and the concatenated sequence, phylogeographic analyses were conducted using maximum likelihood (ML) tree inference and haplotype network estimation. An ML tree was inferred using IQ-TREE v. 2.2.2.6 (Minh et al., 2020). The downloaded target sequences of *B. oreas*, *B. toni* (outgroup), and *B. barbatula* (outgroup) were added for each tree inference.

Partitions were set for each codon position, and the nucleotide substitution model selected using ModelFinder (Kalyaanamoorthy et al., 2017) based on the Bayesian Information Criterion (BIC) was applied to each partition (Table S6). Node support was assessed using the Ultrafast Bootstrap approach (Hoang et al., 2018) with 1,000 replicates. The resulting tree was visualised using interactive Tree of Life (iToL) v. 6 (Letunic and Bork, 2024). A haplotype network was constructed using POPART v. 1.7 (http://popart.otago.ac.nz; Leigh and Bryant, 2015) with the Templeton–Crandall–Sing (TCS) algorithm (Clement et al., 2000). Genetic groups were identified based on the bootstrap value in the ML tree and the 3-step clades derived from the network following the nesting rule (Templeton and Sing, 1993). The geographic distribution of these mtDNA groups was visualised using QGIS v. 3.40 (available at https://qgis.org/en/site/).

### 2.8 Divergence time estimation

To estimate the divergence timing of species and clades, a time-calibrated phylogenetic tree was inferred using a concatenated dataset of ND5 and cyt *b* (Fig. S2). As outgroups, we included 12 Nemacheilidae species, primarily those associated with calibration points. Among these, *L. echigonia* (AB080170) and *Barbatula* sp. Korea (PP279918) lacked ND5 data; however, because they were essential for the analysis, we included them in the dataset with all ND5 sequences set as “N”. We first estimated an ML tree using the same method as described above and examined the resulting topology. For divergence time estimation, we used BEAST v. 2.7.8 (Bouckaert et al., 2019) and estimated the site model for each gene partition using the bModelTest package v. 1.3.3 (Bouckaert and Drummond, 2017) (Table S6). A relaxed clock log-normal model was applied, and the tree prior was set to the Birth-Death model; both were linked across partitions. Two calibration points were applied following Ooyagi et al. (2018) and Šlechtová et al. (2025). The first calibration point was the uplift age of the Central Highlands (3–7 Ma; Machida et al., 2006), used as a prior for the divergence between eastern and western *Lefua echigonia* (log-normal, offset = 0, mean = 5.0, standard deviation = 0.25). The second calibration point was based on the oldest known fossil record of the genus *Triplophysa* (*T. opinata*; 5.3–16.0 Ma) (Prokofiev, 2007). Calibration densities for the fossil record were calculated using CladeAge v2.1.0 (Matschiner et al., 2017), with general parameter values for Osteichthyes: net diversification rate = 0.041–0.081, turnover rate = 0.0011–0.37, and sampling rate = 0.0066–0.01806 (Foote and Miller, 2007; Matschiner et al., 2017; Santini et al., 2009). All other parameters were set to default values. MCMC analyses were run for 100,000,000 generations, with sampling every 10,000 generations. We confirmed that all parameters had effective sample size (ESS) values above 200 using Tracer v1.7.2 (Rambaut et al., 2018). The resulting tree file excluded the first 40% of trees as burn-in and combined the remaining trees into a consensus tree using TreeAnnotator v2.7.7.

### 2.9 Library preparation for quantitative fish eDNA metabarcoding

To estimate the eDNA concentration of all fish species, including *B. oreas*, quantitative fish eDNA metabarcoding analysis was performed with MiFish-U primer set (Miya et al., 2015) and standard DNAs. The paired-end library preparation with a two-step PCR was performed in 12 µL of reaction mixture according to the described method in Tsuji et al. (2022). Briefly, in first-round PCR, the 12S rRNA region of fishes (ca 170 bp) and three internal standard DNAs (5, 25, 50 copies per reaction, respectively: Table S7) were amplified together using MiFish-U primer set. PCR negative controls with ultrapure water instead of the eDNA sample were added in all first PCR runs. The second-round PCR was performed to add index-(Hamady et al., 2008) and adapter sequences for the Illumina sequencing platform. The indexed products of the second-round PCR were pooled, and the target bands (ca. 370 bp) were excised using 2% E-Gel SizeSelect Agarose Gels (Thermo Fisher Scientific, MA, USA). The prepared DNA library was sequenced (2 × 150 bp paired-end sequencing) using illumina NovaSeq 6000. The sequencing process was outsourced to Novogene Japan (Tokyo, Japan) and conducted using the GigaBase sequencing service.

The bioinformatics analysis was performed using the PMiFish pipeline v.2.4.1 (https://github.com/rogotoh/PMiFish). The settings for each parameter are as follows: Rarefaction = No, Length Filter = 120, Depth Filter = 0.01%, Correct_error = Yes, UIdentity = 98.5. Species assignment was performed using a local BLASTN search with the reference database including all inhabiting freshwater fish taxa at study sites (MiFish DB ver. 37) and the sequences of three internal standard DNAs. As all study sites were in freshwater areas, any saltwater fish detected were excluded from subsequent analyses. To obtain a sample-specific standard line, linear regression analysis (lm function in R), was performed using the sequence reads of internal standard DNAs and their known copy numbers (the intercept was set as zero). For each sample, the number of eDNA copies for each taxon was calculated using each sample-specific standard line: eDNA concentration (copies/µL) = the number of sequence reads/regression slope of the sample-specific standard line (Table S8). Ideally, the target 1,000 bp of *B. oreas* should be quantified using the real-time PCR method. However, due to methodological limitations preventing accurate quantification of longer sequences (typically > 200 bp), the concentration of the approximately 170 bp 12S rRNA fragments quantified by the qMiFish method was used in subsequent statistical analyses as the eDNA concentration of *B. oreas*.

## 3 Results

### 3.1 Sequencing of 1,000 bp from eDNA samples and haplotyping

Of the 52 samples analysed, sequencing of approximately 1,000 bp of ND5 and cyt *b* was successful in 34 samples (Table S9, Fig. 2A). Each sample that succeeded sequencing yielded 2,037 ± 1,175 (mean ± standard deviation) reads. The N50 read length was 1,065.9 ± 63.5 bp. 92 ± 9 % of the reads obtained from each sample were mapped to the reference sequence of ND5 and cyt *b*. Of the remaining 18 samples, sequencing was attempted in 6 samples, but no valid data could be obtained. The approximate time required after eDNA extraction was 30 minutes for multiplex PCR, 30 minutes for PCR amplicon concentration and purification, 1 hour for circularisation of DNA fragments, and 2–3 days for full-plasmid DNA sequence analysis (outsourced; GENEWIZ from AZENTA Life Sciences). The financial cost for library preparation and sequencing was 4,390 JPY (≈ 25.55 EUR, ≈ 29.87 USD) per sample (Table S10).

**Figure 2.**
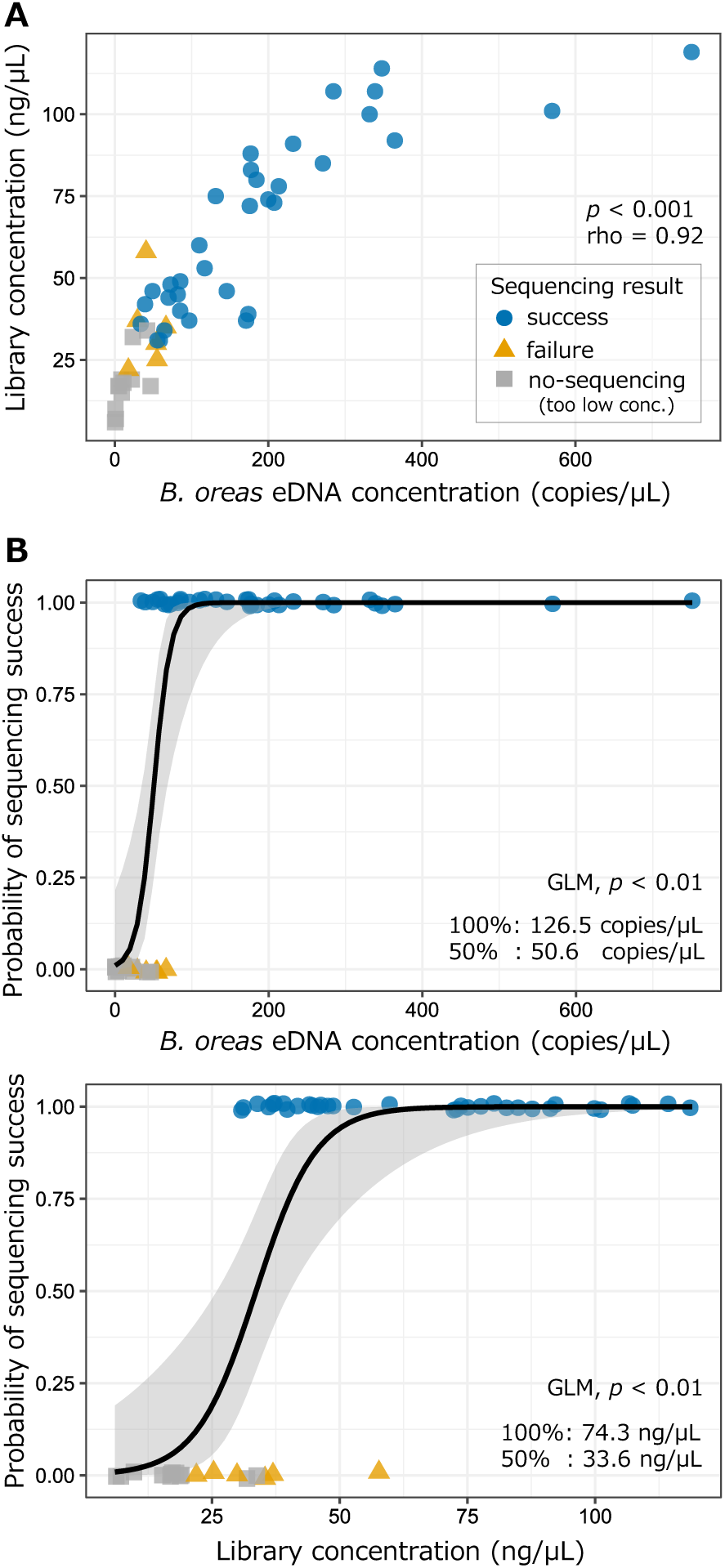
(A) The relationship between *B. oreas* eDNA concentration in a sample (12S rRNA, ca. 170 bp) quantified using the qMiFish metabarcoding approach and the prepared library concentration for full-length plasmid sequencing (Spearman’s rank correlation test, rho = 0.92, *p* < 0.001). Point shapes and colours indicate the result of long-fragment eDNA sequencing. (B) The results of full-length plasmid sequencing as a function of their eDNA concentration in a sample (upper) or the prepared library concentration (lower). The distinction between plot colours and shapes is the same as in (A). Plots are vertically staggered to minimise overlapping. The solid lines (Y-axis) indicate probabilities of sequencing success, respectively (GLM, *p* < 0.01, both panels). The shaded area in both panels corresponds to the 95% confidence intervals

A strong positive correlation was observed between the *B.oreas* eDNA concentration (MiFish region, 12S rRNA, ca.170 bp) in the samples before PCR and the concentration of the final sequencing library for full-plasmid DNA sequence analysis (Spearman’s rank correlation test, rho = 0.92, *p* < 0.001; Fig. 2A). Furthermore, GLM analysis revealed that the probability of successful sequencing was significantly related with the concentration of *B.oreas* eDNA in the sample (z = 2.84, *p* < 0.01) and the concentration of the prepared sequence library (z = 3.04, *p* < 0.01), respectively (Fig. 2B). Based on the obtained relationship, to achieve a 50% and 100% probability of successful sequencing, it was estimated that eDNA concentrations of 50.6 copies/µL and 126.5 copies/µL, and library concentrations of 33.6 ng/µL and 74.3 ng/µL are required, respectively.

When assigning read frequencies to unique haplotypes under different ploidy assumptions (diploid to hexaploid), the diploid assumption showed the highest frequency for all samples with two or more SNPs (Table S11). However, in some samples, the frequency of assigned reads was similar between diploid and triploid assumptions. For these samples, SNPs were visually confirmed using IGV, and the triploid assumption was adopted for ND5 in st.25 and cyt *b* in st.11, st.14, st.18, st.25 (Table S11). The number of detected haplotypes per sample was 1.76 ± 0.50 for ND5 and 1.75 ± 0.67 for cyt *b*. The number of detected haplotypes was consistent between ND5 and cyt *b* in most samples, although some samples showed an additional haplotype in one of the regions. In total, 53 haplotypes were identified in ND5 and 52 haplotypes in cyt *b* from 34 samples. Furthermore, for each sample, the haplotypes of ND5 and cyt *b* detected were concatenated considering genetic groups and frequencies, resulting in 62 concatenated sequences (Table S5).

### 3.2 Phylogenetic relationships and phylogeographic patterns

In all haplotype data sets—ND5, cyt *b* and their concatenated sequence—two highly differentiated clades (Clade A and Clade B) were identified in *B. oreas* (Fig. 3, S3, S4). In all haplotype data sets, Clade A and Clade B were distributed separately in the southern and northern areas of Hokkaido but were detected sympatrically at three sites (st.13, st.14 and st.15; Fig. 3, S3, S4). Each clade comprised five (Clade A) and three (Clade B) regional subgroups. The haplotype network formed a bottleneck-type characterised by deep divergences between clades and groups, along with multiple missing haplotypes (Fig. 3B). The A5 group within Clade A corresponds to the previously reported group 1 (Yatsuyanagi et al. 2024), while the remaining groups, A1, A2, A3 and A4, had a distribution pattern that further subdivided the previously reported two groups (groups 2 and 3). Clade B was divided into three subgroups, consistent with previous studies. However, the distribution ranges of each group shown in this study did not correspond with those previously reported (Yatsuyanagi et al. 2024), and the distribution areas were relatively clearly distinguished between the eastern and western parts of northern Hokkaido. Interestingly, as in previous studies, sequences obtained from Sakhalin specimens were classified into the B3 group. Furthermore, the collection site information for all NCBI-registered sequences was consistent with the distribution areas of the corresponding regional subgroups identified by the eDNA analysis (Table S2, Fig. 3, S4).

**Figure 3.**
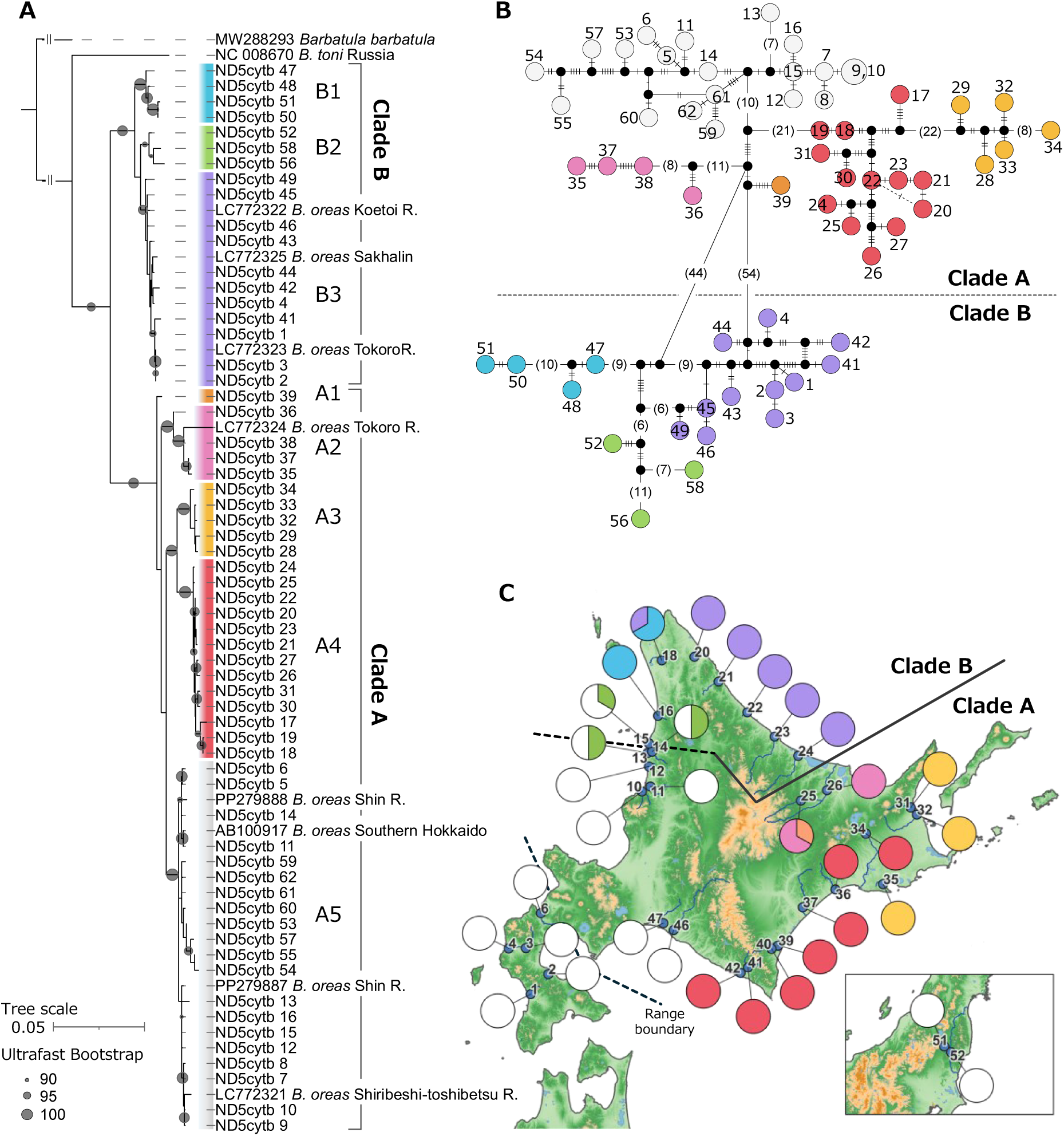
Genetic population structure of *B. oreas* inferred using 1,968 bp of concatenated sequence (ND5, 953 bp and cyt *b*, 1,015 bp) sequences from long-fragment eDNA analysis. The colour scheme used for each regional subgroup is consistent across all panels. (A) Maximum likelihood phylogenetic tree. The ID indicates the detected haplotype (Table S5) and the accession number of the downloaded sequences. Ultrafast bootstrap support values (> 90%) are indicated by circle sizes. (B) Haplotype network. Short bars and numbers in parentheses indicate the number of base differences between neighbouring haplotypes. Black small circles indicate missing haplotypes. (C) Distribution map of each regional subgroup. Pie charts represent the relative frequencies of detected groups

**Figure 4.**
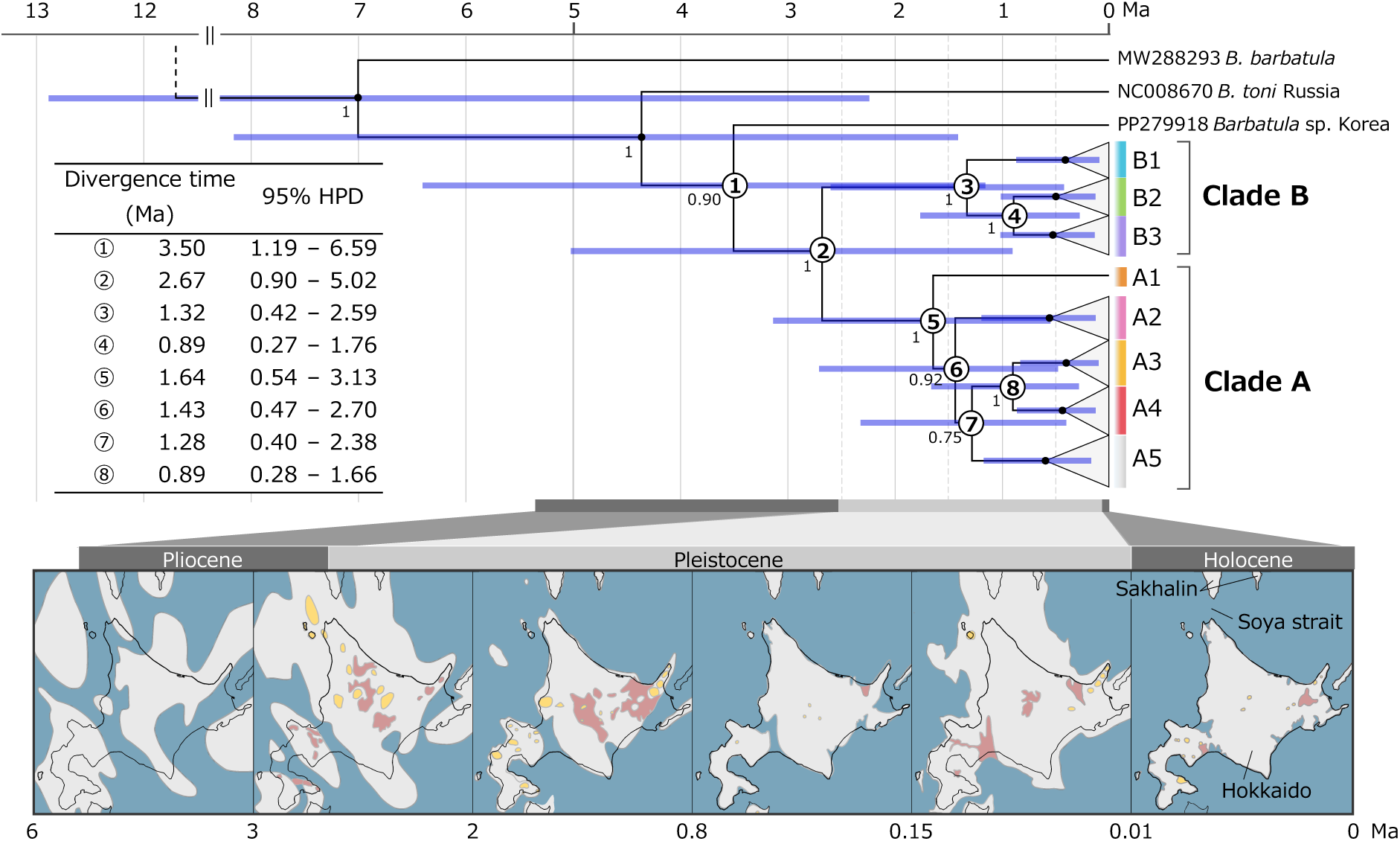
Time-calibrated Bayesian phylogenetic tree based on 1,968 bp of concatenated sequences (ND5, 953 bp and cyt *b*, 1,015 bp) showing estimated divergence times among *Barbatula* species (see Fig. S2 for the overall view). Numbers below the node point indicate Bayesian posterior probabilities (BPP). Purple node bars represent the 95% highest posterior density (HPD) of divergence time. The divergence time and 95% HPD for major divergences ①–⑧ were shown in the table. The map below the phylogenetic tree shows the approximate distribution of land (grey) and sea (blue) around Hokkaido and Sakhalin for each geological period (modified from Minato, 1973). Red and yellow areas indicate the range of volcanic ash and pumice ejected as pyroclastic flows, and the distribution of lava and volcanic ash, respectively.

In both ND5 and cyt *b* haplotype data sets, AMOVA indicated a significant genetic differentiation between the two clades (ND5, variance explained = 81.9%, *Φ*CT = 0.82, *p* < 0.05; cyt *b*, variance explained = 76.4%, *Φ*CT = 0.13, *p* < 0.05). Also, a significant genetic differentiation among subgroups (ND5, variance explained = 15.8%, *Φ*SC = 0.88, p < 0.001; cyt *b*, variance explained = 22.0%, *Φ*SC = 0.93, *p* < 0.001: Table 1).

**Table 1.** Summary of the Analysis of Molecular Variance (AMOVA) Results.

| Region | Source of variation | SSD | MSD | DF | Variance explained % | p-value | Phi-statistics |
| --- | --- | --- | --- | --- | --- | --- | --- |
| ND5 | Between two clades | 1.68.E-02 | 1.68.E-02 | 1 | 81.9 | < 0.05 | 0.82 |
|  | Among subgroups | 4.85.E-03 | 8.09.E-04 | 6 | 15.8 | < 0.001 | 0.88 |
|  | Error | 9.18.E-04 | 2.04.E-05 | 45 | 2.2 | - | 0.97 |
|  | Total | 2.26.E-02 | 4.34.E-04 | 52 |  | - | - |
| cyt b | Between two clades | 1.72.E-02 | 1.72.E-02 | 1 | 76.4 | < 0.05 | 0.76 |
|  | Among subgroups | 7.06.E-03 | 1.18.E-03 | 6 | 22.0 | < 0.001 | 0.93 |
|  | Error | 6.73.E-04 | 1.53.E-05 | 44 | 1.7 | - | 0.98 |
|  | Total | 2.49.E-02 | 4.88.E-04 | 51 |  | - | - |

### 3.3 Divergence times inferred from concatenated sequences

Using all 62 concatenated sequences, the divergence time was estimated based on the partitioned ND5 and cyt *b* regions (Fig. 4, S2). The sister species to *B. oreas* was estimated to be Barbatula sp. Korian population; their divergence time was estimated at 3.50 Ma [95% highest posterior density (HPD), 1.19–6.59]. The time to the most recent common ancestor (tMRCA) of *B. oreas* was estimated at 2.67 Ma (0.90–5.02). The tMRCA of Clade A was estimated at 1.64 Ma (0.54– 3.13), and the divergence between Clade A2 and A3 + A4 + A5 was estimated at 1.43 Ma (0.47– 2.70). Clade A5 and A3 + A4 were estimated to have diverged 1.28 Ma (0.40–2.38), and subsequently, Clade A3 and A4 diverged 0.89 Ma (0.28–1.66). The tMRCA of Clade B was estimated at 1.32 Ma (0.42–2.59), and the divergence between Clade B2 and B3 was estimated at 0.89 Ma (0.27–1.76).

### 3.4 Quantitative fish eDNA metabarcoding

Sequencing of the 55 libraries yielded 336,624 ±40,252 (mean ± SD) raw reads per library (52 field samples, one filt-NC and two PCR-NC; Table S12). A total of 62 fish taxa were detected and quantified after bioinformatics analysis (Table S13). The R^2^ values of the sample-specific standard line ranged from 0.70 to 1.00. The regression slope ranged from 68.81 to 2,948.11 (Table S8). *Barbatulus oreas* was detected in all samples, and the DNA copy number was estimated. The number of copies in each detection sample of *B. oreas* and the top 10 native species (*Oncorhynchus masou masou, Tribolodon hakonensis, Salvelinus* sp./*Parahucho perryi, Gymnogobius* sp., *Tribolodon sachalinensis, Oncorhynchus* sp., *Tridentiger* sp., *Pungitius* sp., *Rhinogobius* sp., *Cottus amblystomopsis*) with the highest number of detection samples among other detected species (i.e., those likely to coexist with *B. oreas*) are shown in Fig. S5. Among them, DNA was detected at concentrations exceeding the thresholds of 50% (50.6 copies/µL) or 100% (126.5 copies/µL) sequencing success of 1,000 bp in multiple species.

## 4 Discussion

In this study, we developed a simple and cost-effective long-fragment eDNA analysis method for evaluating intraspecific genetic variation and successfully sequenced approximately 1,000 bp of two mtDNA regions from eDNA samples. By reconstructing longer sequence data than those used in conventional eDNA analyses, we revealed the population structure of *B. oreas* at a higher resolution. The capability to evaluate genetic structure at a practical resolution — without the need for extensive field capture sampling, complex laboratory workflows, and high analysis costs — represents a significant advance for genetic diversity surveys by improving the feasibility of such surveys and accelerating their social implementation.

### 4.1 Requirements and practical feasibility of long-fragment eDNA sequencing

Our results demonstrated that sequencing success strongly depended on both the concentration of target eDNA and the prepared library, providing empirical thresholds that can guide the practical use of the method. This finding is significant for field studies, as it offers clear criteria for sample collection and quality assessment before sequencing. It is widely recognised that eDNA is fragmented in the environment due to enzymatic degradation, microbial activity, and physical processes (Barnes and Turner, 2016; Eichmiller et al., 2016; Strickler et al., 2015; Tsuji et al., 2017); however, recent studies suggest that detection potential can be enhanced by applying protocols optimised for detecting longer eDNA fragments (West and Deagle, 2025). The thresholds for successful sequencing of approximately 1,000 bp estimated in this study were within the range of eDNA concentrations typically observed for common species under natural conditions (50% success, 50.6 copies/µL; 100% success, 126.5 copies/µL; note that this refers to the concentration of fragments approximately 170 bp in length, not 1000 bp) (e.g., aquatic plants, Kodama et al., 2021; fish, Saito et al., 2022; shrimp, Wu et al., 2019). This interpretation was further supported by the fact that several other species, not targeted in our sampling design, were also detected above the thresholds in multiple samples (Fig. S5). These results suggested that obtaining approximately 1,000 bp amplicons is not restricted to exceptionally high-quality samples such as tank water or high-density habitat water but can be achieved under realistic field conditions. Furthermore, even when the initial eDNA concentration is below the threshold, the required library concentration (50% success, 33.6 ng/µL; 100% success, 74.3 ng/µL) might still be achieved by applying enrichment steps during library preparation. For example, in this study, DNA fragments from four PCR replicates were pooled per sample, but increasing the number of replicates within a realistic range according to the abundance of the target species (e.g., up to eight; Doi et al., 2021; Ficetola et al., 2015) could further enhance the applicability of long eDNA sequencing. These insights provide a basis for refining protocols and expanding the range of ecological and conservation studies that can benefit from long eDNA sequencing.

### 4.2 A practical balance of resolution, labour, and financial cost

Our proposed method reduces both technical and financial barriers for high-resolution eDNA analysis and facilitates its broader application in ecological and conservation studies across academia, governments, industry, and the private sector. This study focused on proposing a method that can be performed even without abundant funding, technical expertise, or experimental resources by utilising affordable commercial sequencing services, thereby bypassing the initial setup process for long-read analysis environments. The workflow of sequencing library preparation and bioinformatic analysis involves only basic laboratory procedures and analysis tools, allowing for rapid turnaround within several days (Fig. 1B, C). The per-sample cost of library preparation and sequencing was approximately 4,390 JPY (≈ 25.55 EUR, ≈ 29.87 USD; Table S10), significantly lower than probe-based target capture methods and shotgun sequencing (Jensen et al., 2021). Furthermore, when a long-read sequencer is already available in the laboratory, amplified eDNA fragments can be sequenced directly by adding adapter and tag sequences without circularising them (cf. Maggini et al., 2024b). In this case, the analysis cost per sample can be further reduced.

The approximately 1,000 bp targeted in this study corresponds to the sequence length commonly employed in mtDNA-based population genetics and phylogeography studies using tissue DNA (e.g., Chiba, 1999; Fuke et al., 2024; Tojo et al., 2017). Short-read eDNA metabarcoding, which has been primarily used to date, is inexpensive and simple; however, its critical limitation has been insufficient analytical resolution due to being restricted to short fragments (approximately ≤ 400 bp; Nakajima and Tsuri, 2024; Sigsgaard et al., 2020; Tsuji et al., 2025a; Turon et al., 2020; Wakimura et al., 2023; Yatsuyanagi et al., 2024). By overcoming this limitation, our proposed method—coupled with the inherent advantages of eDNA analysis, such as simplified field surveys and the ability to analyse multiple species simultaneously—will further elevate the applicability and value of eDNA-based genetic diversity assessments.

Moreover, it is known that longer DNA fragments exist in the environment, making it possible to target sequences longer than 1,000 bp (Doorenspleet et al., 2025; Maggini et al., 2024b; West and Deagle, 2025). However, there is a trade-off between eDNA fragment length and detectability, with an inherent risk of false negatives (West and Deagle, 2025). Therefore, in this study, using samples with eDNA concentrations typically found in environmental samples, 1,000 bp was targeted for analysis as the length enabling practical resolution. On the other hand, when higher-concentration eDNA samples are available, analysing longer sequences is expected to yield higher-resolution, current-state information on genetic diversity (Doorenspleet et al., 2025).

4.3 *High-resolution phylogeographic structure of* B. oreas

By obtaining approximately 1,000 base pairs of sequence from each of two mtDNA regions, two major clades and multiple regional subgroups were clearly identified, revealing a higher-resolution population genetic structure of *B. oreas*. The previous study based on shorter eDNA sequences also reported two clearly differentiated clades and multiple regional subgroups (Yatsuyanagi et al., 2024). However, due to the short length of the analysed sequences, boundaries among regional subgroups were unclear, leading to some groups being conflated or overlooked. In this study, the ability to analyse longer sequences enabled higher-precision identification of regional subgroup boundaries. Specifically, within Clade B, three regional subgroups—previously suggested to have complex distributions—were confirmed to be clearly spatially separated. Furthermore, sister groups A3 and A4, previously considered a single group, were clearly identified as distinct groups through analysis based on longer sequences. The longer sequence length also enabled the application of AMOVA, confirming significant differentiation between the two clades and among regional subgroups, revealing a clear and deep population genetic structure in *B. oreas*. Interestingly, while the three freshwater fish species *B. oreas*, *Rhynchocypris percnurus sachalinensis*, and *Lefua nikkonis* are known to have historically migrated from Sakhalin to Hokkaido via a land bridge formed across the Soya Strait, their genetic structure patterns show almost no correspondence (Goto, 1978; Ooyagi et al., 2018). This suggests that despite sharing the same expansion route, the three species likely differed in their timing of migration, patterns of distribution expansion, and demographic population dynamics.

Results from divergence time estimates provided new insights into the migration process of *B. oreas* into Hokkaido and its subsequent distribution expansion. This study showed that two clades of *B. oreas* diverged approximately 2.67 Ma, a period coinciding with the disappearance of the land bridge connecting Hokkaido and Sakhalin (3.0–2.0 Ma, Fig.4; Minato, 1973).

Furthermore, consistent with the previous study, group B3 includes haplotypes from Sakhalin specimens, suggesting that Clade B maintained genetic exchange with the Sakhalin population relatively recently (Yatsuyanagi et al., 2024). Considering these findings, it is inferred that the common ancestor first dispersed via Sakhalin to Hokkaido, and subsequent marine transgressions geographically isolated the two regions, leading to the divergence of Clade A in Hokkaido.

Furthermore, during the Late Pleistocene, significant sea-level drops led to the reformation of a land bridge across the Soya Strait, allowing Clade B or their ancestral population to migrate southward into Hokkaido (Fig. 4). This scenario is also supported by the fact that no clear geographic barrier exists at the distribution boundary between the Clade A and B. Furthermore, at the distribution boundary between the two Clades, they were detected sympatrically in multiple rivers, suggesting the existence of a secondary contact zone. However, all studies to date have been based on mtDNA, and nuclear genome information has not been considered. Additionally, the current survey sites are limited to lower reaches, leaving the population structure in inland regions unexplored. Future studies combining comprehensive sampling, including inland areas, with nuclear genome analysis to clarify the presence of reproductive isolation between clades and details of demographic population dynamics will provide deeper insights into the mechanisms of species differentiation and distribution history.

The distinct genetic differentiation and distribution patterns of *B. oreas* suggest a deep relationship between past geological/climatic events and the processes shaping biodiversity in Hokkaido. This study indicates that the regional subgroups belonging to Clade A diverged sequentially between approximately 1.64 and 0.89 Ma (Fig.4), coinciding with a period of heightened volcanic activity in Hokkaido (2.0–0.8 Ma; Geology of Japan “Hokkaido Region” Editorial Committee, 1990; Minato, 1973). Furthermore, parts of the distribution ranges of groups A3 and A5 are thought to have been temporarily submerged during this period. This submersion likely caused habitat fragmentation and isolation, which in turn may have promoted population differentiations. Indeed, the haplotype network indicates structures suggesting bottlenecks (Fig. 3B), interpreted as small populations surviving in refugia and subsequently re-expanding their distributions. Such evolutionary processes have also been reported in other aquatic species inhabiting Hokkaido (e.g., *Hynobius retardatus*, Azuma et al., 2013; *Cambaroides japonicus*, Koizumi et al., 2012), supporting the maintenance of populations in limited locations under harsh environmental conditions. On the other hand, while diversification within Clade B likely occurred in the geographically isolated Sakhalin, this study could not identify the specific factors driving it. Due to the lack of specimens from Sakhalin, further sampling, including Sakhalin, is necessary to clarify this issue.

Detailed phylogeographic information not only clarifies the history of population formation but also contributes to estimating the origin and pathways of recent anthropogenic introductions, as well as assessing the extent of genetic disturbance. Our results indicate that individuals originating from the southern Hokkaido region have been introduced into multiple non-native areas. *Barbatula oreas* are popular as ornamental fish among dedicated aquarists, leading to the capture of individuals in their habitats and their sale nationwide. In southern Hokkaido, the extreme concentration of people (Statistics Bureau Ministry of Internal Affairs and Communications, 2020), combined with developed transportation infrastructure, including New Chitose Airport (CTS; serving approximately 20 million passengers annually; https://x.gd/9dYnu), makes the transport of individuals to other regions relatively easy. Given these circumstances, the risk of individual being transported by private collectors and aquarium retailers from southern Hokkaido is considered higher than in other areas. On the other hand, within the native range, it has been informally suggested by aquaculture practitioners that *B. oreas* might have been unintentionally introduced through salmon release. Nevertheless, each regional subgroup is generally detected within its respective localities, suggesting that large-scale genetic disruption is unlikely. This is likely attributable to the localised implementation of salmon stocking programs, in which the egg collection, rearing, and release of individuals are largely conducted within the same region, with minimal interregional transfer (Yamao and Amano, 2019). The increasing ease of movement of people and goods over the past few decades has heightened the risk of introducing alien species both domestically and internationally (Dukes et al., 1999; Seebens et al., 2021). Detailed phylogeographic information is essential for estimating the origin and migration routes of introduced populations, and it would provide the scientific basis for risk assessment and the development of effective management strategies to prevent the invasion and spread of alien species.

### 4.4 Broader applicability and limitations of long-fragment eDNA sequencing

The applicability of long-*fragment* eDNA sequencing is not confined to *B. oreas*, but can be extended to a wide range of taxa and ecosystems. By using multiple or group-specific primers, the method can be applied not only to the simultaneous analysis of multiple regions, as shown here, but also to multi-species surveys, offering a promising direction for future applications. A key requirement for expanding the applicability of this method is the design of species- or group-specific primers. This has also occasionally been cited as a barrier to species-specific eDNA detection using real-time PCR and electrophoresis (Freeland, 2017; Sahu et al., 2025). However, more than 17 years have passed since eDNA detection of macroorganisms was first reported, and knowledge and expertise regarding eDNA primer design should have steadily accumulated. Furthermore, when applying metabarcoding techniques, it is possible to remove non-targeted sequences if a well-curated reference database is available (Tsuji et al., 2023a; Weitemier et al., 2021). Therefore, primer design remains an important consideration, but it is unlikely to be a decisive barrier.

PCR-based eDNA approaches are affected by amplification biases, false negatives due to low concentration or degraded DNA, and false positives due to error amplification (Evans and Lamberti, 2018; Goldberg et al., 2016; Rishan et al., 2023). In particular, when detecting long eDNA, eDNA released from individuals that were nearby at the time of sampling and had not yet undergone degradation is likely to be amplified more easily (Doorenspleet et al., 2025). Previous studies based on short eDNA fragments have demonstrated positive correlations between estimated and true haplotype (allele) frequencies (Andres et al., 2021; Marshall and Stepien, 2019; Nakajima and Tsuri, 2024; Tsuji et al., 2020b), but it remains to be examined whether similar patterns can be confirmed for long eDNA, ideally through validation against capture-based data.

Furthermore, controlling false negatives and false positives while retaining true haplotype and allele diversity is a central challenge in metabarcoding-based analyses of genetic variation, exemplifying the classical trade-off between type I (false positive) and type II (false negative) errors (Couton et al., 2023; Macé et al., 2022; Turon et al., 2020). Since this study focused on phylogenetic analysis, we emphasised minimising false positives by applying a relatively high VAF threshold (20%), thereby restricting detection to reliable haplotypes with a high likelihood of existing. Consequently, it is likely that rare haplotypes were overlooked, as was also suggested by the relatively low haplotype diversity observed at each site. However, if high-frequency haplotypes were detected at each site, a few false negatives would have little impact on the estimation of major phylogenetic groups and their geographic patterns (Pedersen et al., 2021; Tsuji et al., 2023a). On the other hand, when aiming to detect genetic differentiation at a more local scale, setting a lower VAF threshold to reduce false negatives could be an option. In this case, the risk of false positives simultaneously increases. Nevertheless, since frequency-based summary statistics, such as *Φ*ST, in population genetics analysis are relatively robust to low-frequency variants, the impact of some false positives should likely be limited (Couton et al., 2023; Nakajima and Tsuri, 2024). Error control remains a critical challenge in eDNA-based assessments of genetic diversity; however, adjusting thresholds in line with specific research objectives would offer a practical solution to address this issue.

The main limitation of our proposed method is its low applicability to nuclear DNA markers (single-copy region). While mtDNA is easily detected from eDNA samples due to its high copy number, the detection of nuclear DNA from eDNA samples remains more challenging because of its comparatively low copy number (generally about 1/10 or less of mtDNA; Pinfield et al., 2019; Saito et al., 2023; Tsuji and Shibata, 2025). Nevertheless, due to characteristics such as maternal inheritance and the absence of recombination, the limitations of inferring and discussing population structure based solely on mtDNA are widely recognised (Ballard and Whitlock, 2004; Galtier et al., 2009). On the other hand, even if 1 kbp of the nuclear region can be sequenced, the highly conserved nature of nuclear sequences often limits the number of polymorphic sites, leaving challenges for improving the resolution of population analysis.

Despite its limited application due to technical complexity, labour intensity, high financial cost and the need for advanced prior knowledge of genetic variants, probe-based capture methods and shotgun sequencing have demonstrated potential for detecting nuclear DNA from eDNA samples (Jensen et al., 2021; Liu et al., 2024; Manel et al., 2025). PCR amplification of microsatellite loci may also hold promise, but challenges remain regarding concentration, species-specific detection, and the availability of reference data (Andres et al., 2021, 2023). Although breakthroughs will be required, it will be essential to advance the continued development and integration of multiple approaches—including the long-fragment eDNA sequencing method developed in this study—to achieve more straightforward and reliable analyses of nuclear genetic diversity from eDNA in the near future.

## Authors’ contributions

S.T. conceived and designed the research. S.T. and N.S. performed a field survey. S.T. performed molecular experiments. All authors conducted data analysis and visualisation. S.T. wrote the early draft and completed it with significant input from all authors.

## Data accessibility

All raw sequences were deposited in the DDBJ Sequence Read Archive with the accession numbers: PRJDB37702 (PSUB043335), DRR753625-DRR753679.

## Benefit-Sharing Statement

Benefits from this research accrue from the sharing of our data and results on public databases as described above.

## Acknowledgements

We thank K. Watanabe (Kyoto University) for valuable advice; Y. Katayama and Y. Miuchi (Kyoto University) for providing materials (tissue sample and fish picture); and H. Doi (Kyoto University) for laboratory facilities and partial support for consumables. This work was supported by JSPS KAKENHI Grant Number 23K13967 and Management Expenses Grants for National Universities.

---

Figure S1. Conceptual diagram showing how haplotypes of ND5 and cyt *b* were concatenated for each sample. Haplotypes connected by lines represent sequence pairs used for concatenation. These pairs were selected based on the detection frequency of each haplotype and their regional subgroup, prioritising combinations that were as consistent as possible

Figure S2. Time-calibrated Bayesian phylogenetic tree based on 1,968 bp of concatenated sequences (ND5, 953 bp and cyt *b*, 1,015 bp) showing estimated divergence times among Barbatula species. The first calibration point (CP1) was the uplift age of the Central Highlands (3–7 Ma; Machida et al., 2006), used as a prior for the divergence between eastern and western *Lefua echigonia* (log-normal, offset = 0, mean = 5.0, standard deviation = 0.25). The second calibration point (CP2) was based on the oldest known fossil record of the genus *Triplophysa* (*T. opinata*; 5.3–16.0 Ma) (Prokofiev, 2007).

Figure S3. Genetic population structure of *B. oreas* inferred using 953 bp of ND5 sequences from long-fragment eDNA analysis. The colour scheme used for each regional subgroup is consistent across all panels. (A) Maximum likelihood phylogenetic tree (Haplotype ID; Table S4), with downloaded sequences (accession No. in NCBI). Ultrafast bootstrap support values (> 90%) are indicated by circle sizes. (B) Haplotype network. Short bars and numbers in parentheses indicate the number of base differences between neighbouring haplotypes. Black small circles indicate missing haplotypes. (C) Distribution map of each regional subgroup. Pie charts represent the relative frequencies of detected groups

Figure S4. Genetic population structure of *B. oreas* inferred using 1,015 bp of cyt *b* sequences from long-fragment eDNA analysis. The colour scheme used for each regional subgroup is consistent across all panels. (A) Maximum likelihood phylogenetic tree (Haplotype ID; Table S4), with downloaded sequences (accession No. in NCBI). Ultrafast bootstrap support values (> 90%) are indicated by circle sizes. (B) Haplotype network. Short bars and numbers in parentheses indicate the number of base differences between neighbouring haplotypes. Black small circles indicate missing haplotypes. (C) Distribution map of each regional subgroup. Pie charts represent the relative frequencies of detected groups

Figure S5. The top 10 native freshwater fish species frequently detected alongside *B. oreas* in qMiFish analysis. Numbers above each species indicate the number of positive detections out of 52 samples. The plot shows the distribution of estimated eDNA concentrations (log₁₀ copies/µL) for each species in samples where they were detected. Solid and dashed horizontal lines represent the eDNA concentrations at which full-length plasmid sequencing success rates reach 100% and 50%, respectively

