## Supplemental figure for "A Practical and Cost-Effective Approach to Long-Fragment eDNA Sequencing for High-Resolution Genetic Diversity Assessment"

| sample | ND5 | | | | cyt *b* | | | | concatenated sequence |
|---|---|---|---|---|---|---|---|---|---|
| | Hap.ID | seq. | freq. | group | Hap.ID | seq. | freq. | group | |
| 1 | ND5_H1 | ATGC | 0.8 | A1 | cytb_H1 | TCAG | 0.9 | A1 | ATGCTCAG |
| 1 | ND5_H2 | ATTC | 0.2 | A1 | cytb_H2 | TCGA | 0.1 | A1 | ATTCTCGA |
| 2 | ND5_H1 | ATGC | 1.0 | A1 | cytb_H1 | TCAG | 0.6 | A1 | ATGCTCAG |
| 2 | | | | | cytb_H3 | TAGA | 0.4 | A1 | ATGCTAGA |
| 3 | ND5_H1 | ATGC | 1.0 | A1 | cytb_H1 | TCAG | 0.7 | A1 | ATGCTCAG |
| 3 | | | | | cytb_H4 | CTGA | 0.3 | A2 | |
| 4 | ND5_H1 | ATGC | 0.5 | A1 | cytb_H1 | TCAG | 0.7 | A1 | ATGCTCAG |
| 4 | ND5_H2 | ATTC | 0.2 | A1 | | | | | ATTCTCAG |
| 4 | ND5_H3 | ACCG | 0.3 | A2 | cytb_H4 | CTGA | 0.3 | A2 | ACCGCTGA |

Fig. S1

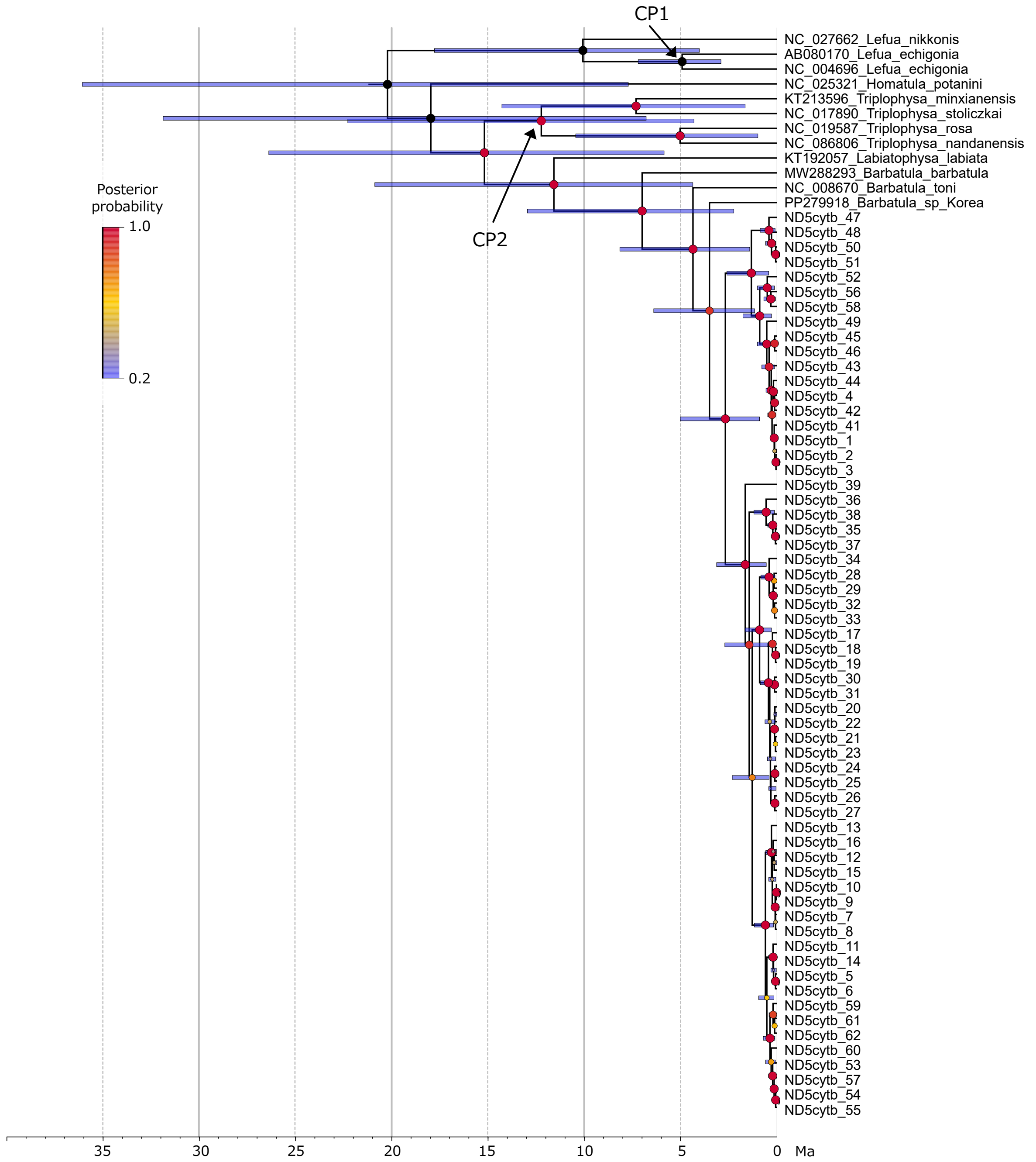
CP1
CP2
Posterior probability
1.0
0.2
NC_027662_Lefua_nikkonis
AB080170_Lefua_echigonia
NC_004696_Lefua_echigonia
NC_025321_Homatula_potanini
KT213596_Triplophysa_minxianensis
NC_017890_Triplophysa_stoliczkai
NC_019587_Triplophysa_rosa
NC_086806_Triplophysa_nandanensis
KT192057_Labiatophysa_labiata
MW288293_Barbatula_barbatula
NC_008670_Barbatula_toni
PP279918_Barbatula_sp_Korea
ND5cytb_47
ND5cytb_48
ND5cytb_50
ND5cytb_51
ND5cytb_52
ND5cytb_56
ND5cytb_58
ND5cytb_49
ND5cytb_45
ND5cytb_46
ND5cytb_43
ND5cytb_44
ND5cytb_4
ND5cytb_42
ND5cytb_41
ND5cytb_1
ND5cytb_2
ND5cytb_3
ND5cytb_39
ND5cytb_36
ND5cytb_38
ND5cytb_35
ND5cytb_37
ND5cytb_34
ND5cytb_28
ND5cytb_29
ND5cytb_32
ND5cytb_33
ND5cytb_17
ND5cytb_18
ND5cytb_19
ND5cytb_30
ND5cytb_31
ND5cytb_20
ND5cytb_22
ND5cytb_21
ND5cytb_23
ND5cytb_24
ND5cytb_25
ND5cytb_26
ND5cytb_27
ND5cytb_13
ND5cytb_16
ND5cytb_12
ND5cytb_15
ND5cytb_10
ND5cytb_9
ND5cytb_7
ND5cytb_8
ND5cytb_11
ND5cytb_14
ND5cytb_5
ND5cytb_6
ND5cytb_59
ND5cytb_61
ND5cytb_62
ND5cytb_60
ND5cytb_53
ND5cytb_57
ND5cytb_54
ND5cytb_55
35
30
25
20
15
10
5
0
Ma

Fig. S2

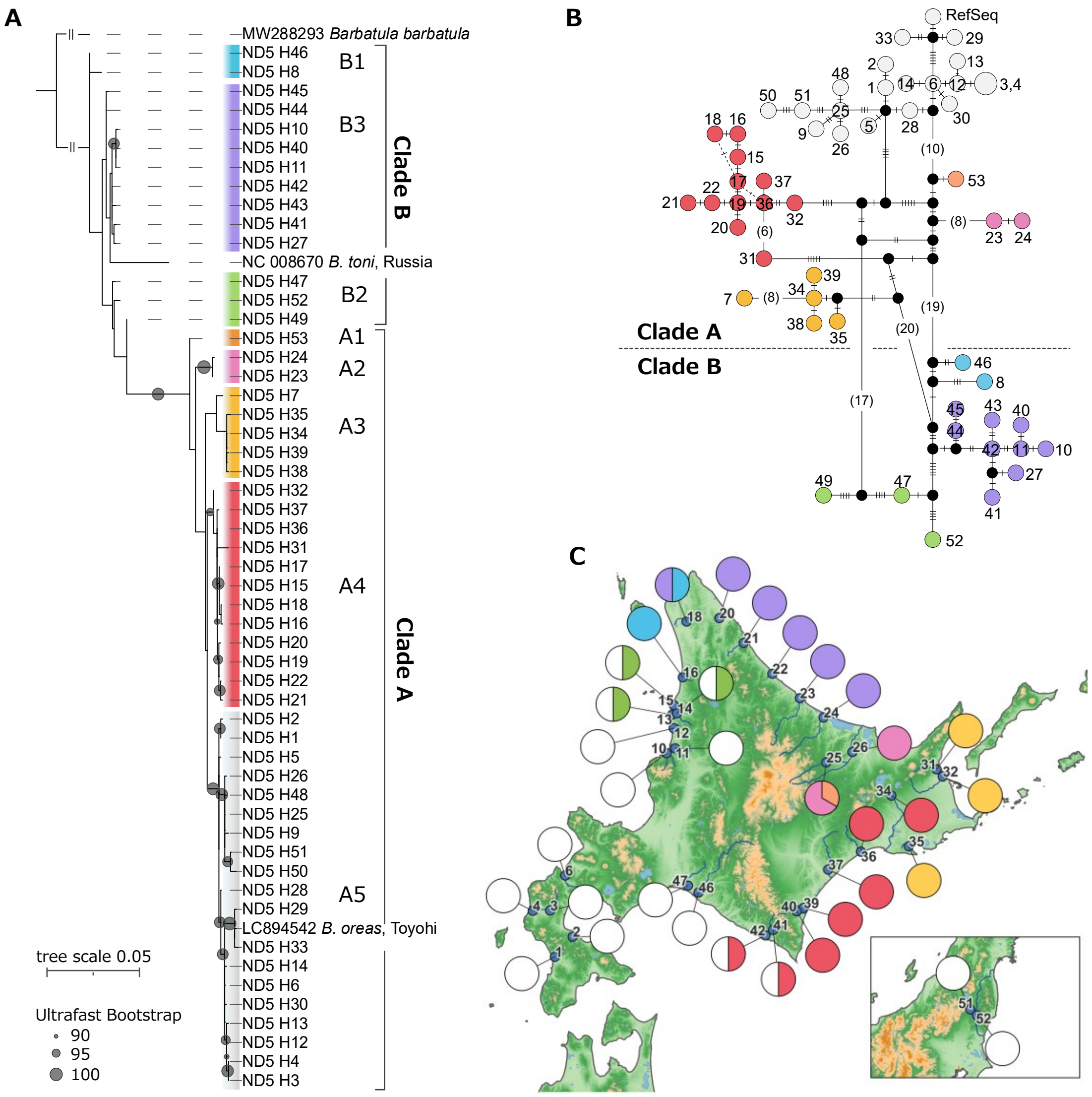

A
MW288293 *Barbatula barbatula*
ND5 H46
ND5 H8
B1
ND5 H45
ND5 H44
ND5 H10
ND5 H40
ND5 H11
ND5 H42
ND5 H43
ND5 H41
ND5 H27
B3
Clade B
NC 008670 *B. toni*, Russia
ND5 H47
ND5 H52
ND5 H49
B2
ND5 H53
A1
ND5 H24
ND5 H23
A2
ND5 H7
ND5 H35
ND5 H34
ND5 H39
ND5 H38
A3
ND5 H32
ND5 H37
ND5 H36
ND5 H31
ND5 H17
ND5 H15
ND5 H18
ND5 H16
ND5 H20
ND5 H19
ND5 H22
ND5 H21
A4
Clade A
ND5 H2
ND5 H1
ND5 H5
ND5 H26
ND5 H48
ND5 H25
ND5 H9
ND5 H51
ND5 H50
ND5 H28
ND5 H29
LC894542 *B. oreas*, Toyohi
ND5 H33
ND5 H14
ND5 H6
ND5 H30
ND5 H13
ND5 H12
ND5 H4
ND5 H3
A5
tree scale 0.05
Ultrafast Bootstrap
90
95
100
B
RefSeq
Clade A
Clade B
C


Fig. S3

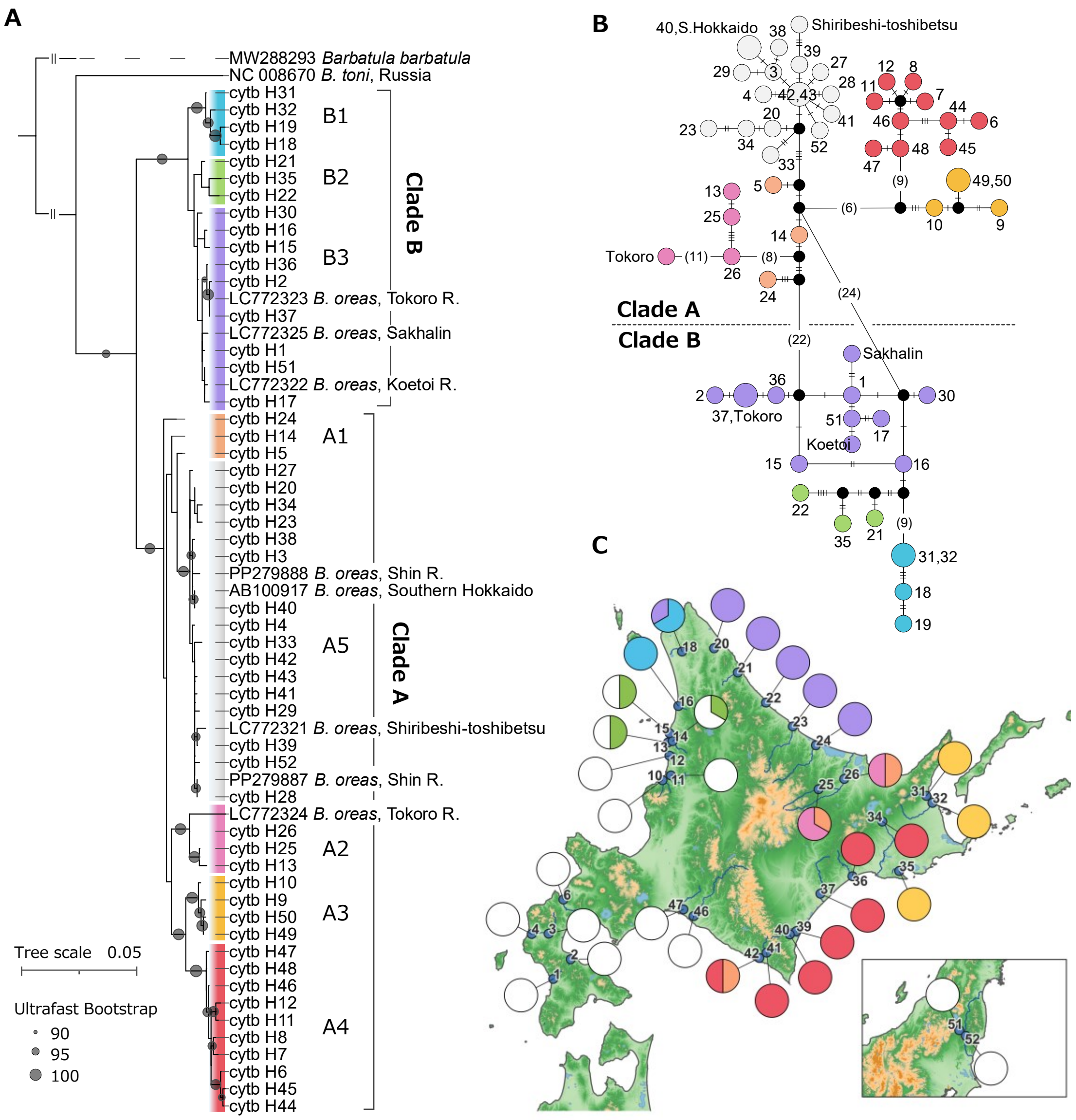
A
MW288293 Barbatula barbatula
NC 008670 B. toni, Russia
cytb H31
cytb H32
cytb H19
cytb H18
cytb H21
cytb H35
cytb H22
cytb H30
cytb H16
cytb H15
cytb H36
cytb H2
LC772323 B. oreas, Tokoro R.
cytb H37
LC772325 B. oreas, Sakhalin
cytb H1
cytb H51
LC772322 B. oreas, Koetoi R.
cytb H17
cytb H24
cytb H14
cytb H5
cytb H27
cytb H20
cytb H34
cytb H23
cytb H38
cytb H3
PP279888 B. oreas, Shin R.
AB100917 B. oreas, Southern Hokkaido
cytb H40
cytb H4
cytb H33
cytb H42
cytb H43
cytb H41
cytb H29
LC772321 B. oreas, Shiribeshi-toshibetsu
cytb H39
cytb H52
PP279887 B. oreas, Shin R.
cytb H28
LC772324 B. oreas, Tokoro R.
cytb H26
cytb H25
cytb H13
cytb H10
cytb H9
cytb H50
cytb H49
cytb H47
cytb H48
cytb H46
cytb H12
cytb H11
cytb H8
cytb H7
cytb H6
cytb H45
cytb H44
B1
B2
B3
Clade B
A1
A5
Clade A
A2
A3
A4
Tree scale 0.05
Ultrafast Bootstrap
90
95
100
B
40,S.Hokkaido
Shiribeshi-toshibetsu
42,43
Tokoro
Clade A
Clade B
Sakhalin
37,Tokoro
Koetoi
49,50
31,32
C

Fig. S4

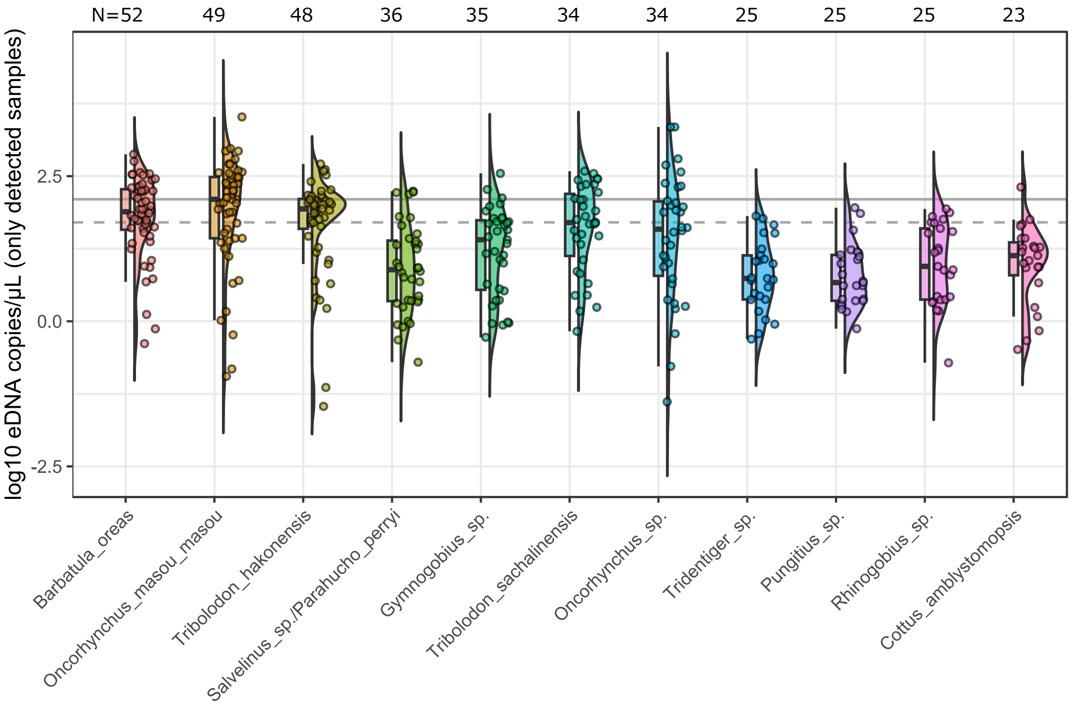

N=52
49
48
36
35
34
34
25
25
25
23
log10 eDNA copies/μL (only detected samples)
2.5
0.0
-2.5
Barbatula_oreas
Oncorhynchus_masou_masou
Tribolodon_hakonensis
Salvelinus_sp./Parahucho_perryi
Gymnogobius_sp.
Tribolodon_sachalinensis
Oncorhynchus_sp.
Tridentiger_sp.
Pungitius_sp.
Rhinogobius_sp.
Cottus_amblystomopsis


Fig. S5
